# Optimizing for generalization in the decoding of internally generated activity in the hippocampus

**DOI:** 10.1101/066670

**Authors:** Matthijs A.A. van der Meer, Alyssa A. Carey, Youki Tanaka

## Abstract

The decoding of a sensory or motor variable from neural activity benefits from a known ground truth against which decoding performance can be compared. In contrast, the decoding of covert, cognitive neural activity, such as occurs in memory recall or planning, typically cannot be compared to a known ground truth. As a result, it is unclear how decoders of such internally generated activity should be configured in practice. We suggest that if the true code for covert activity is unknown, decoders should be optimized for generalization performance using cross-validation. Using ensemble recording data from hippocampal place cells, we show that this cross-validation approach results in different decoding error, different optimal decoding parameters, and different distributions of error across the decoded variable space. In addition, we show that a minor modification to the commonly used Bayesian decoding procedure, which enables the use of spike density functions, results in substantially lower decoding errors. These results have implications for the interpretation of covert neural activity, and suggest easy-to-implement changes to commonly used procedures across domains, with applications to hippocampal place cells in particular.

## Introduction

The decoding of neural activity is a powerful and ubiquitous approach to understanding information processing in the brain. Decoding is typically cast as a mapping from neural data to a sensory or motor variable, such as the identity of a visually presented object or the reaching direction of a motor action; the same idea can be applied to more abstract or even hidden states such as context or past history. By comparing a decoded (“reconstructed”) variable with the actual value, the contributions of features such as spike timing, adaptation, and correlations to decoding accuracy can be quantified (Nirenberg and Latham, 2003; Panzeri et al., 2015; Schneidman, 2016). Based on the nature and accuracy of the decoder output under various conditions, inferences may be drawn about the possible functions of neural populations carrying such signals and the circuitry responsible for generating them (Georgopoulos et al., 1986; Bialek et al., 1991; Pillow et al., 2008). These decoding approaches share the property that when a known stimulus value is available along with neural data, decoding performance can be optimized relative to a known “ground truth”(i.e. the actual stimulus value).

Increasingly so, however, decoding is also applied to brain activity occurring in the *absence of any overt stimulus or action* (Georgopoulos et al., 1989; Johnson et al., 2009; King and Dehaene, 2014). Such internally generated activity occurs, for instance, during processes such as planning, deliberation, visual imagery and perspective-taking, memory recall and sleep. A well-studied example is provided by studies of hippocampal activity recorded in rodents, which exhibits internally generated sequences of neural activity that appear to depict behavioral trajectories during sleep and wakeful rest (“replay”; Skaggs and McNaughton 1996; Nadasdy et al. 1999; Davidson et al. 2009; Pfeiffer and Foster 2013), and during the theta rhythm while task-engaged (“theta sequences”, Foster and Wilson 2007; Gupta et al. 2012; Chadwick et al. 2015). Replay is thought to reflect an off-line consolidation process from a fast-learning, episodic-like short-term memory trace in the hippocampus into a semantic-like neocortical knowledge structure (McClelland et al., 1995; Káli and Dayan, 2004; Girardeau et al., 2009; Carr et al., 2011), but also plays a role in on-line task performance, and can depict trajectories that are not well explained by consolidation processes, such as those towards a behaviorally relevant goal and never-experienced paths (O’Neill et al., 2006; Jadhav et al., 2012; Dragoi and Tonegawa, 2013; Ólafsdóttir et al., 2015). Theta sequences may enable one-shot learning, and/or play a role in on-line prediction during behavior (Lisman and Redish, 2009; Malhotra et al., 2012; Feng et al., 2015).

How should we interpret the *content* of such internally generated activity? The intuition that replay has a clear resemblance to activity observed during active behavior can be formalized by simply applying the same decoder used to decode activity during overt behavior (Tatsuno et al., 2006; Kloosterman, 2012; Shirer et al., 2012). However, in the rodent hippocampus there are also obvious differences between the two types of activity, such as the compressed timescale and different instantaneous population firing rates (Skaggs and McNaughton, 1996; Lee and Wilson, 2002; Buzsáki, 2015). More generally, there is now overwhelming evidence that hippocampal “place cells” are better viewed as encoding many possible stimulus dimensions rather than just place; these may include relatively low-level properties such as running speed, information about objects and events, and complex history– and context-dependence (Huxter et al., 2003; Lin et al., 2005; McKenzie et al., 2014; Allen et al., 2016). Thus, it is unlikely that the mapping between neural activity and encoded location (the “encoding model”) remains the same between overt and covert epochs, raising the possibility of biases in our ability to decode specific stimulus values, such as different positions along a track.

To address the above issues, we provide several practical improvements to commonly used decoding procedures, of particular use for applications to internally generated activity. In acknowledgment of the likely different encoding model in force during overt and covert neural activity, we emphasize that decoding performance should be optimized for generalization performance (i.e. to do well on withheld data not used to estimate the parameters of the decoder). We compare different splits of the data, and show that these not only result in different overall decoding accuracy, but also in different accuracy distributions over the stimulus space. In particular, these nonuniformities (biases) in accuracy only become apparent when optimizing for generalization to data not included in the training set. Because decoding internally generated activity also involves such generalization, the interpretation of decoding such activity should be informed by the known shape of this bias. Finally, we show that regardless of the type of split used, decoding accuracy can be improved by relaxing the assumption of integer spike counts used in the common Bayesian decoding procedure (Zhang et al., 1998; Johnson and Redish, 2007; Pfeiffer and Foster, 2013).

## Materials and Methods

### Overview

Our aim is to describe how the output of decoding hippocampal ensemble activity depends on the configuration of the decoder. In particular, we examine two components: (1) the split between training and testing data, and (2) the parameters associated with the estimation of firing rates and tuning curves (the encoding model). Both are described in the *Analysis* section. All analyses are performed on multiple single unit data recorded from rats performing a T-maze task, described in the *Behavior* section. Data acquisition, annotation, and pre-processing steps are described in the *Neural data* section.

All preprocessing and analysis code is publicly available on our GitHub repository,https://github.com/vandermeerlab/papers. Data files are available from our lab server on request by e-mail to the corresponding author.

### Neural data

#### Subjects and overall timeline

Four male Long-Evans rats (Charles River and Harlan Laboratories), weighing 439-501 g at the start of the experiment, were first introduced to the behavioral apparatus (described below; 3-11 days) before being implanted with an electrode array targeting the CA1 area of the dorsal hippocampus (details below). Following recovery (4-9 days) rats were reintroduced to the maze until they ran proficiently (0-3 days), at which point daily recording sessions began. On alternate days, rats were water-or food-restricted. In parallel with the maze task, some rats (R042, R044, R050) were trained on a simple Pavlovian conditioning task in a separate room (data not analyzed).

#### Behavioral task

The apparatus was an elevated T-maze, constructed from wood, painted matte black with white stripes applied to the left arm (Figure 1) and placed on a metal frame approximately 35 cm in height. The distance from the start of the central stem to the ends of the arms was 272 cm (R042) or 334 cm (R044, R050, R064; these numbers are subject IDs). 6% sucrose (∼0.1 ml) was dispensed upon reaching the end of the left arm, and food (5 pellets of Test Diet 5TUL 45 mg pellets) was dispensed upon reaching the end of the right arm.

**Figure 1:**
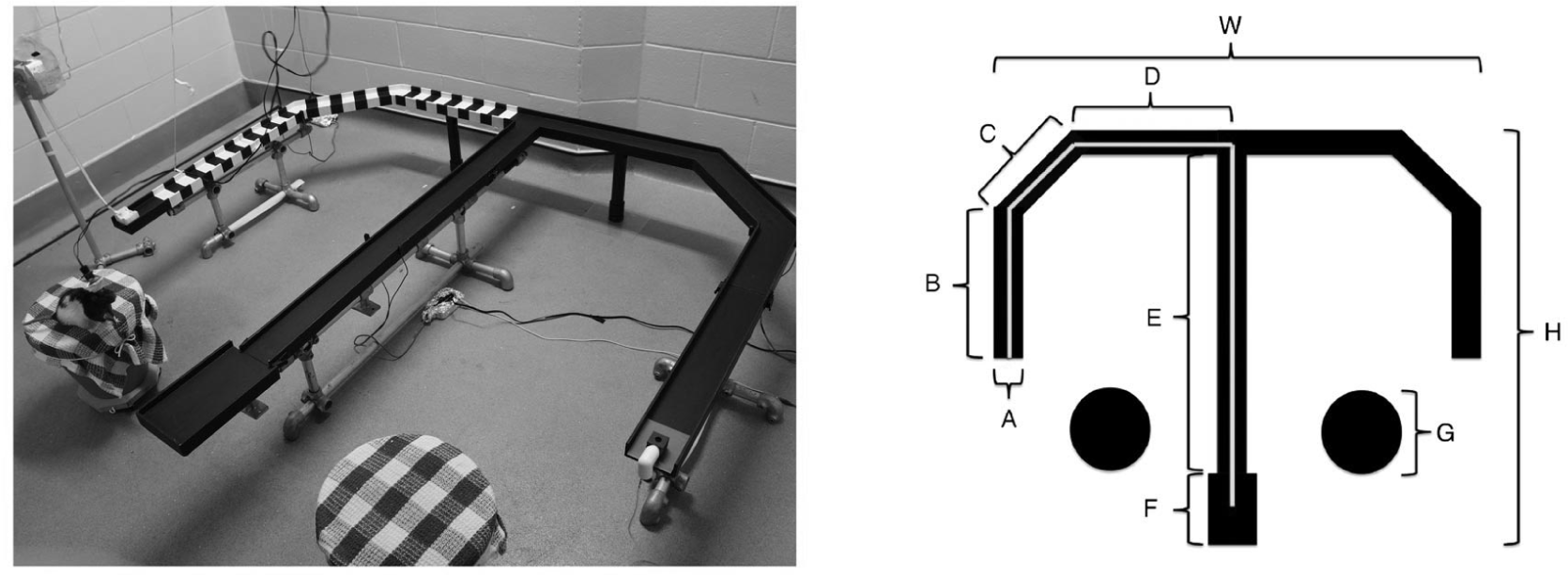
Behavioral apparatus. In daily recording sessions, rats ran approximately 20 trials on an elevated T-maze. Trials were free-choice except for a small number of forced trials in which access to one of the arms was prevented by a barrier to ensure that at least 5 trials for both left and right arms were available for each session. Track dimensions were: width 10 cm (A), total maze height 167 cm (H), total maze width 185 cm (W), total path length 334 cm (white trajectory, R042 excepted, who had a shorter B segment for a total path length of 285 cm).

Daily recording sessions consisted of (1) a pre-behavior recording epoch, taken as the animal rested on a recording pedestal (terracotta pot lined with towels; 20-30 min), (2) approximately 20 trials on the maze, with an intertrial interval (30-240 s) on the recording pedestal after each trial, and (3) a post-behavior recording epoch (10-20 min). A trial was defined as a run from the starting point at the base of the central stem to one of the reward locations; photobeams at the track ends were used to find pairs of crossings defining the shortest interval between leaving the base and arriving at an end. Only data from runs on the track were analyzed here.

Because rats were food– or water-restricted, they tended to prefer choosing the arm leading to the reward to which their access was limited. On some sessions, access to a preferred arm was blocked with a movable barrier to ensure sampling of the non-preferred arm (forced choice). Trials on which the animal turned around, or exhibited other disruptive behaviors (climbing on the barrier, extended grooming, etc.) were excluded from analysis.

#### Electrode arrays and surgery

Subjects were each implanted with a single-bundle microelectrode array targeting the CA1 region of dorsal hippocampus in the right hemisphere (AP −4.0mm, ML +2.5mm). R042 and R044 were each implanted with a 15-tetrode 1-reference array, and R050 and R064 were each implanted with a 16-tetrode 4-reference array. Surgical procedures were as described previously (Malhotra et al., 2015). Briefly, the skull was exposed and a ground screw was placed through the contralateral parietal bone. Arrays were lowered to the surface of the cortex through a craniotomy, and the remaining exposed opening was sealed with a silicone polymer (KwikSil). Then, the arrays were anchored to the skull using small screws and acrylic cement. Rats were given a minimum recovery period of four days, during which antibiotics and analgesics were administered, before retraining began. Tetrodes were slowly advanced to the CA1 layer over a period of 4-9 days. The first recording sessions began no sooner than nine days after surgery. All procedures were performed in accordance with the Canadian Council for Animal Care (CCAC) guidelines, and pre-approved by the University of Waterloo Animal Care Committee (protocol 10-06).

#### Recording methods

Neural activity from all tetrodes and references was recorded on a Neuralynx Digital Lynx SX data acquisition system using HS-36-LED analog buffering headstages tethered to a motorized commutator. Local field potentials, filtered between 1-425 Hz, were continuously sampled at 2 kHz. Spike waveforms, filtered between 600-6000 Hz, were sampled at 32 kHz for 1 ms when the voltage exceeded an experimenter-set threshold (typically 40-50µ V) and stored for offline sorting. Acquired signals for all rats (except R042, whose data was recorded relative to animal ground) were referenced to an electrode located in the corpus callosum, dorsal to the target recording site. A video tracking algorithm recorded the rat’s position based on headstage LEDs picked up by an overhead camera, sampling at 30 Hz. All position data was linearized by mapping each 2-dimensional position sample onto the nearest point of an ideal linearized trajectory on the track, drawn for each session by the experimenter. Position samples further than 25cm from this idealized trajectory were treated as missing values.

#### Preprocessing and annotation

Signals were preprocessed to exclude intervals with chewing artifacts and high-amplitude noise transients where necessary. All spiking data was initially clustered into putative units automatically (KlustaKwik, K. D. Harris) and then manually checked and sorted (MClust 3.5, A. D. Redish). Highly unstable units and units that fired fewer than 100 spikes in a recording session were excluded. Recording locations were histologically confirmed to lie in the dorsal CA1 cell layer. A total of 2017 units were recorded from 4 rats across 24 sessions (Table 1); 889 of these were units were rated as questionable isolation quality by the experimenter and kept separate for later analysis.

**Table 1:** Total neural units for each rat across each of their six recording sessions. Numbers inparentheses indicate how many of the numbers listed were units rated as questionable. Sessions listed in italics were excluded due to insufficient number of recorded units.

| Rat ID | Session 1 | Session 2 | Session 3 | Session 4 | Session 5 | Session 6 | Total |
| --- | --- | --- | --- | --- | --- | --- | --- |
| **R042** | 22 (*13*) | 74 (39) | 107 (40) | 64 (43) | 73 (40) | 59 (31) | **399 (206)** |
| **R044** | *17 (11)* | *13 (9)* | *43 (21)* | 53 (26) | 50 (27) | *41 (22)* | **217 (116)** |
| **R050** | 72 (42) | 94 (40) | 72 (28) | 113 (44) | 128 (36) | 112 (46) | **591 (236)** |
| **R064** | 121 (59) | 136 (47) | 116 (52) | 162 (45) | 178 (68) | 151 (60) | **864 (331)** |

#### Inclusion criteria

Recording sessions with at least 20 units firing a minimum of 25 spikes during “run” epochs (used for tuning curve estimation, described below) for both left and right trials separately were included for analysis. This left out five sessions (four from R044, one from R042) resulting in a total of 19 sessions eligible for analysis.

### Analysis

#### Overview

Our main approach is to employ a standard memoryless Bayesian decoder, common to all analyses and described below. We will vary first, the nature of different splits in the data between “training” and “testing”, and second, parameters associated with the estimation of input firing rates (spike density functions) and input tuning curves (the “encoding model”). In all these cases, the output of the decoding procedure is, for each time bin, a probability distribution over (linearized) position, given the observed spiking activity.

#### Bayesian decoding

We use the canonical Bayesian decoder (Brown et al., 1998; Zhang et al., 1998), specifically the one-step, “memoryless” version with a uniform spatial prior. This procedure (reviewed in detail elsewhere; Johnson et al. 2009; van der Meer et al. 2010; Kloosterman et al. 2014), along with the key parameters varied in this study, is illustrated in Figure 2. The decoded location 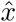 for a given time bin we took to be the mode of the posterior (location with the highest probability; maximum a posteriori). A decoding error can then be defined as the distance to the true position 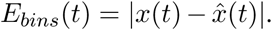. Because *x* has the unit of bins, this quantity is converted into a worst-case error in centimeters as follows: 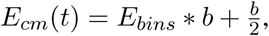, where *b* is the bin size in cm (we used 3 cm for the results reported here, and a time bin τ= 25 ms). Both the estimation of tuning curves (the encoding model, described below) and the decoding of spike data were restricted to data when the animal was running (≥5 cm/s).

**Figure 2:**
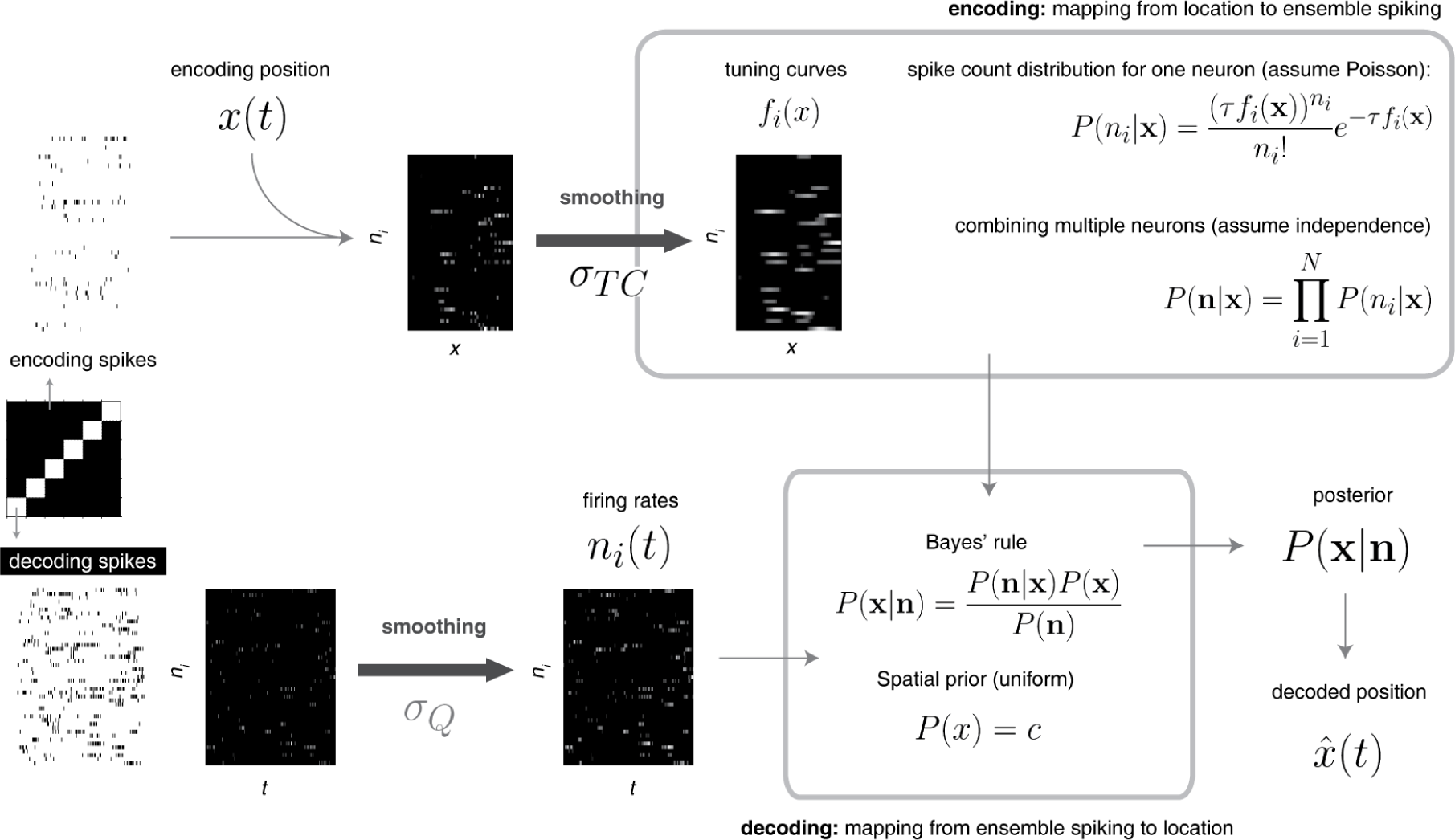
Schematic of the Bayesian decoding scheme. The overall workflow follows the canonical procedure based on the common assumptions of Poisson-distributed spike counts around mean firing rates given by stable tuning curves, and independence between neurons. Crucial variables in the results reported here are (1) the split in the data between trials used for estimating tuning curves (“encoding spikes”) and trials used for decoding (“decoding spikes”; see Figure 3 for a detailed explanation), (2) the width of the Gaussian kernel *σ_TC_* used to smooth the tuning curves (the empirically determined mapping from location to firing rate for each recorded neuron), and (3) the width of the Gaussian kernel *σ_Q_* used to obtain the spike density functions used as the input to the decoder.

We use this decoding procedure here because it has become the *de facto* standard in the hippocampal place cell literature (Kloosterman et al., 2014; Silva et al., 2015; Grosmark and Buzsaki, 2016); however, the manipulations in the present study (discussed below) are general and can be straightforwardly applied to other decoding methods such as optimal linear decoding, regression-based methods and general-purpose classifiers such as support vector machines, et cetera (Pereira et al., 2009; Pillow et al., 2011; Deng et al., 2015).

#### Cross-validation

The data used for the estimation of the encoding model (tuning curves; “training data”) may be the same as the data used for decoding and error estimation (“testing data”), but this need not be the case (Figure 3). We systematically compare different splits between training and testing data, focusing on three specific cases: same-trial decoding (decode each individual trial based on tuning curves obtained from that same trial; Figure 3A), next-trial decoding (decode each individual trial based on tuning curves from the *next* trial; Figure 3C) and leave-one-out decoding (decode each trial based on tuning curves from all trials except the one being decoded; Figure 3D). Decoding errors reported are always for a specific split and this will be reported in the text; note that for all splits used here, each trial is decoded separately, using tuning curves obtained from a set of encoding trials specific to the trial being decoded (this is unlike all-to-all decoding, Figure 3B, in which the same set of all encoding trials is used for every decoding trial). Left and right trials were always treated separately, i.e. only left trials are used to decode left trials, and the same for right trials.

**Figure 3:**
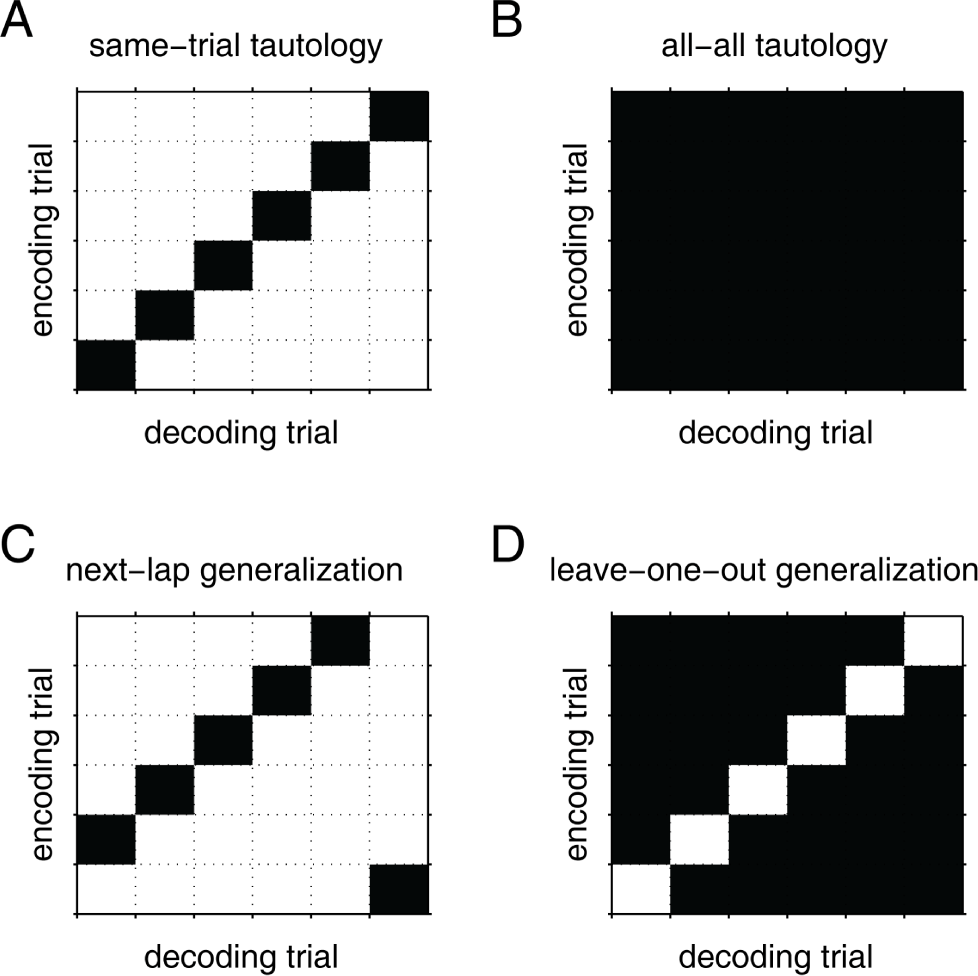
Schematic of different splits between data used for estimating the encoding model (tuning curves, “training data”) and data used for evaluating decoding accuracy (“testing data”). Data splits in the **top row** are “tautological” in that tuning curves are estimated on the same data used for decoding. In contrast, data splits on the **bottom row** measure generalization performance (cross-validation) in the sense that the decoding data was not included in the data used for estimating the encoding model. **Black** cells in the matrices shown indicate trials used to estimate the encoding model. Thus, for instance, the left column in **C** shows that to decode trial 1, tuning curves were estimated from trial 2.

#### Firing rate estimation

Strictly speaking, Bayesian decoding based on the assumption that firing rates are Poisson-distributed requires integer spike counts for the estimation of *P (s|x)* (Figure 2). However, this means that there will be effects of binning, which will become more prominent as the time window (bin) size τ becomes smaller. For instance, if bins only contain 0 or 1 spike, then which side of a bin edge a spike falls on can potentially have a large effect. This issue is prominent in many aspects of spike train analysis, and is typically addressed by convolving the raw spike train to obtain a *spike density function (SDF)*, an estimate of firing rate which varies continuously in time (Cunningham et al., 2009; Kass et al., 2014).

To make the standard Bayesian decoding equations compatible with non-integer spike density functions, we note that the denominator *n_i_* ! does not depend on *x* and can therefore be absorbed into a normalization constant *C* which guarantees that 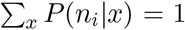 (Eq. 36 in Zhang et al. 1998). For the results presented here, we obtain spike density functions by convolving raw spike trains with a Gaussian kernel with SD *σ_Q_*, discretized at a resolution of 25 ms (the in τ Figure 2).

A possible side effect of using this procedure on the decoding spikes only (i.e. not on the spikes used to estimate the tuning curves, described below) is that firing rate-stimulus combinations that are inconsistent across the ensemble become more likely, e.g. for every individual location *x_i_* in space, there is at least one neuron that assigns *P (x_i_|n)* = 0 (such cases result in the white areas in Figure 4; only sessions in which at least 80% of samples could be decoded were included, except when indicated explicitly in the text). This can be avoided by simply convolving all spikes with the same kernel *σ_Q_*; here we did not do so in order to show the effects of convolving the decoding spikes independently of the encoding model estimation. Smoothing the tuning curves, as described in the next section, is another effective method of avoiding this issue.

**Figure 4:**
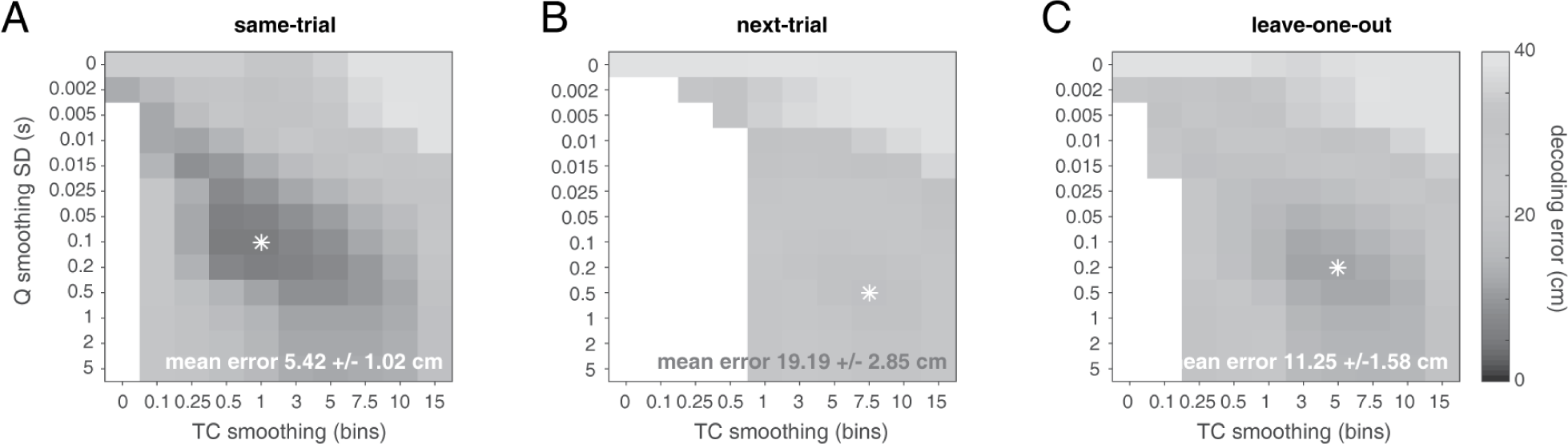
Decoding accuracy for different data splits (**left**: same trial, **middle**: next trial, **right**:leave-one-out) and decoder parameters (vertical axis: standard deviation of Gaussian kernel (in s) used to convolve spike trains, horizontal axis: standard deviation of Gaussian kernel used to convolve tuning curves (in 3 cm bins). Grayscale shows the mean decoding error (in cm) for different parameter and data split combinations. Note that decoding accuracy for some parameter combinations cannot be estimated if temporal smoothing results in decoding spike counts inconsistent with the encoding model (empty data points; see *Methods* for details). Results shown were obtained with a decoding time bin size (τ)of 25 ms; only sessions with at least 20 cells for both left and right trials on the T-maze were included, averaging across left and right trials (19 sessions total). The white asterisk indicates the parameter combination resulting in the lowest mean decoding error; this is the value reported here for each data split, along with the standard error over subjects (n = 4).

#### Encoding model estimation

Bayesian decoding requires an estimate of *P (s|x)*, the probability of observing a firing rate vector *s* for a given stimulus value *x*. As in previous work, we assume firing rates are independent between neurons and Poisson-distributed around some mean rateλ this simplification means that we only need to know the mean firing rate as a function of the stimulus variable, λ(*x*), for each neuron. These are the *tuning curves*, which taken together across all neurons can be thought of as an *encoding model*, i.e. the mapping from stimulus values to neural activity. We estimate tuning curves non-parametrically from the data by (1) restricting the data to intervals when the animal was running on the track (≥5 cm/s; *encoding spikes* in Figure 2), (2) linearizing the position data and binning in bins of 3 cm, (3) obtaining a firing rate histogram by dividing spike count by occupancy for all bins, and (4) optionally smoothing the resulting tuning curve with a Gaussian kernel of standard deviation *σ_TC_* (with units in cm).

#### Inventory of behavioral and neural measures used

Figure 9 shows the distribution, across locations, of a number of behavioral and neural measures which we relate to decoding accuracy. We explain how these are computed in turn.

- *Occupancy*(in seconds; time spent at each location on the track) is computed simply by binning video tracking samples and multiplying the sample counts by the length of each video frame (1/30 s).
- *Place fields* are detected based on session tuning curves, when a contiguous area of at least 15cm is associated with a minimum firing rate of 5 Hz, and a mean firing rate of no more than 10 Hz. For each field (contiguous area) the location of the field is taken to be the neuron’s maximum firing rate in in the field.
- *Tuning curve variability across trials* (Figure 9E-F) is obtained by taking the standard deviation, across trials within each session, of single-trial tuning curves.
- *Bootstrapped tuning curve variability* (Figure 9G-H) is computed by generating a distribution of 1000 resampled tuning curves, with each sample using a random 90% of the position and spiking data. Specifically, every spike is assigned to the position sample closest in time. Then, after selecting a random 90% of position samples, those samples and the associated spikes are removed before computing a tuning curve. A measure of tuning curve variability is then obtained by taking the standard deviation across the distribution of resampled tuning curves.
- *Population vector (PV)* correlations are obtained by correlating, for each location on the track, the tuning curve firing rates (across cells) either between tuning curves obtained from each pair of trials (Figure 9I-J) or between tuning curves from a single trial and the complementary tuning curve of all trials except that one (Figure 9K-L).

## Results

We sought to determine how different configurations of the commonly used one-step Bayesian decoder (Brown et al., 1998; Zhang et al., 1998) relate to the decoding accuracy of position based on ensembles of hippocampal place cells. In particular, we applied different splits to the data, partitioning it into “training” data from which tuning curves were estimated, and “testing” data from which decoding accuracy was determined (a type of cross-validation). In addition, we varied parameters associated with the estimation of tuning curves and firing rates (*σ_TC_* and *σ_Q_* in Figure 2).

Our motivation for exploring different data splits is the question of how internally generated sequences (e.g.“replays”) of neural activity can be decoded in a principled manner. For such sequences, the true mapping from neural activity to stimulus space is generally unknown; after all, there is no true stimulus value to which decoded output can be compared. Under these conditions, decoders should be optimized for generalization performance, i.e. performance on “testing” data not used to “train” the decoder. In statistics and machine learning, such cross-validation is routinely used to prevent overfitting (Hawkins, 2004; Alpaydin, 2014). Applied to the problem of decoding covert sequences, this concept suggests that we choose the decoder which performs best on input data from trials not included in the estimation of tuning curves. Thus, we use data from withheld trials as a proxy for internally generated sequences, such that we can estimate how well various decoders are likely to perform on actual covert sequences.

Specifically, applied to decoding neural data collected across a number of repeated trials, as is the case here in rats running a T-maze task (Figure 1), a number of different splits between testing and training data are possible, illustrated in Figure 3. A commonly used approach in the hippocampal place cell literature is to not perform any split at all, i.e. to estimate tuning curves based on the full data set, and use those to decode the full data set (Figure 3B, Johnson and Redish 2007; Karlsson and Frank 2009; Pfeiffer and Foster 2013; Zheng et al. 2016). We refer to this approach as “tautological” because the same data is used for both. It is possible to do this at different levels of granularity, for instance going down to the single trial level by decoding each individual trial based on tuning curves from that trial (Figure 3A), while maintaining the property that the same data is used for tuning curve estimation and decoding.

### Overall effects of different decoding configurations on accuracy

We found that the best outright decoding performance (as quantified by the error relative to true location) was obtained using such tautological decoding. “Same-trial” decoding performed best of all data splits tested (Figure 4A; average decoding error 5.42±1.02 cm for the best-performing parameters; standard error across subjects). However, if the goal is to optimize decoding performance on trials not included in the training set, the picture changes. Decoding using the *next trial* resulted in a decoding error ∼4-fold worse than the same-trial decoding (19.19±2.85 cm; Figure 4B). Leave-one-out decoding was intermediate between these two (11.25±1.58 cm; Figure 4C), a pattern that held across a wide range of decoding parameters (see also Figure 5 for specific comparisons).

**Figure 5:**
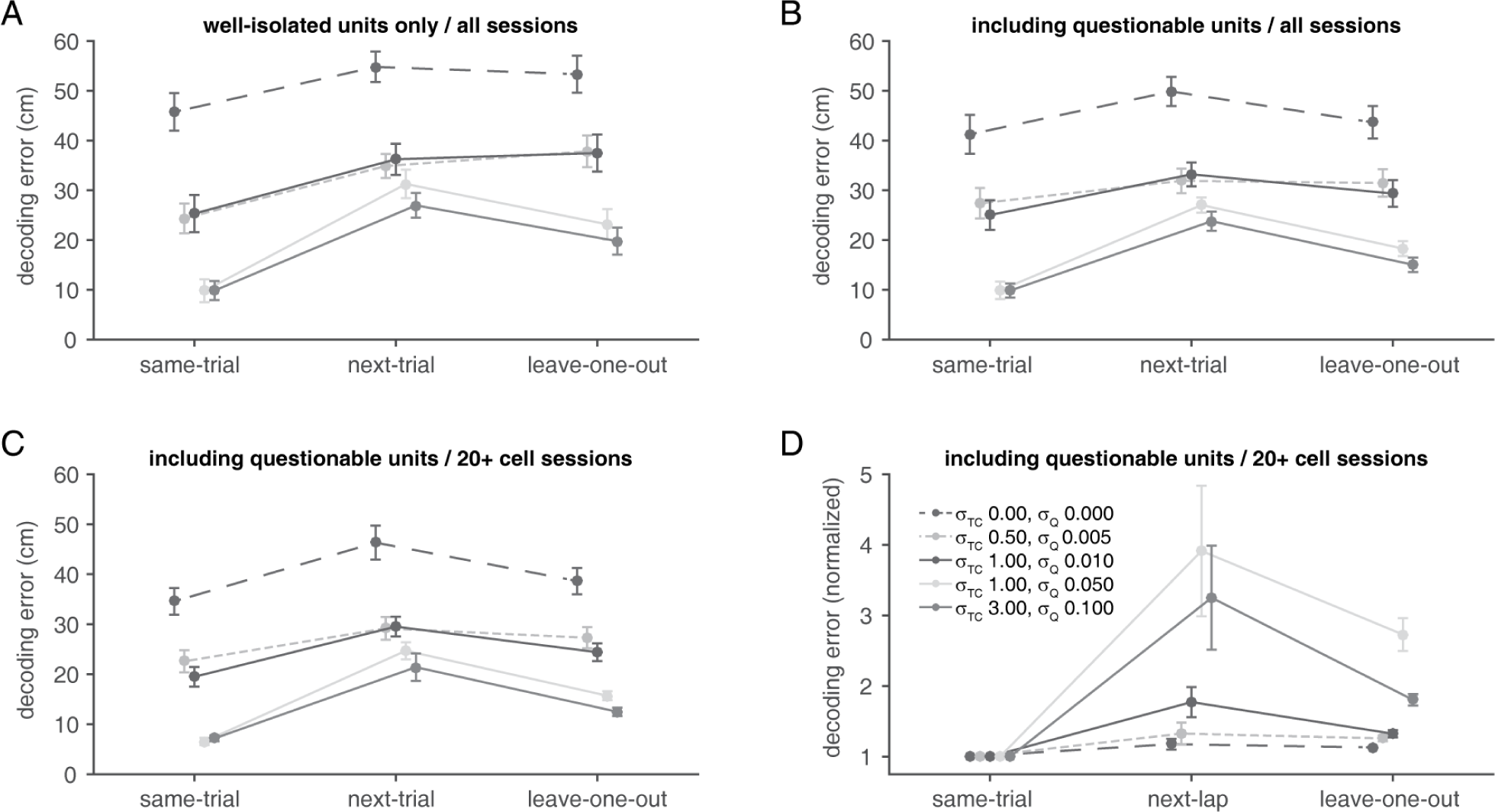
Decoding error for selected parameter and data combinations. Panels **A** and **B** show decoding error run on all sessions (n = 24, i.e. without requiring a minimum number of cells to be active) to compare decoding error when only well-isolated units are used (**A**) or when units of questionable isolation quality are included (**B**). Panels **C** and **D** are replots of the same data as in Figure 4, i.e. for sessions with at least 20 cells that met inclusion criteria (n = 19; see Methods). For panel **D**, decoding error is normalized on a single-session basis to the sa me-trial decoding. Errorbars indicate SEM over subjects (n = 4).

Several other features of Figure 4A-C are worth noting. First, performing no smoothing at all on either the spike trains or the tuning curves (0/0, the data point on the top left of each panel) results in large decoding error. Previous results manipulating the width of the time window indicated minimum error for a time window in the ∼
0.5-1s range (Zhang et al., 1998) this is confirmed here by the error minimum at 0.2 or 0.5 s SD smoothing kernels. Surprisingly however, even very minimal temporal smoothing of the spike trains to be decoded (e.g. a kernel with 5 ms SD) can result in substantial improvements in decoding performance compared to no temporal smoothing (up to 2-fold; see Figure 5 for a close-up of this effect). Second, best decoding accuracy almost invariably required some smoothing of the tuning curves, even when the leave-one-out procedure ensured many trials were used for tuning curve estimation. Third, the parameters yielding optimal decoding accuracy differed between data splits; note for instance how the dark gray area (corresponding to low decoding error) is shifted towards the top left for Figure 4A compared to Figure 4C. Thus, different data splits interact with decoding parameters to produce overall decoding accuracy.

To show more clearly the data in Figure 4 for selected parameter combinations of interest, we plotted separately the raw decoding error (Figure 5A-C) and decoding error normalized to same-trial decoding within each recording session (Figure 5D). Including units with questionable isolation quality decreased decoding error across all conditions (compare Figure 5A-B; see Methods and Table 1 for unit counts), and we therefore used the full set of units including questionable units for all other analyses. Regardless of the set of units used, however, Figure 5 illustrates clearly the large improvement in decoding accuracy of very minimal smoothing (e.g. light dashed line, 5 ms kernel) compared to no smoothing (dark dashed line). Also evident is the performance improvement of the leave-one-out data split over the next-trial data split; this improvement was particularly large for larger smoothing (for which, in turn, overall decoding accuracy was better), for no or minimal smoothing, next-trial and leave-one-out decoding tended not to differ.

### Effect of trial numbers on decoding accuracy

Given that leave-one-out decoding performed as well or better than next-trial decoding, we can ask how this effect depends on the number of trials included in the leave-one-out procedure. This can be of practical importance in determining the number of trials of behavioral sampling will be sufficient for decoding during internally generated activity. Leave-one-out and next-trial decoding can be seen as opposite ends of a spectrum along which the number of trials used to estimate tuning curves is systematically varied. Overall, decoding performance increased as more trials were included, with diminishing returns for larger numbers of trials (Figure 6). As expected from the results in the previous sections, these overall performance gains in absolute and relative decoding accuracy depended on the amount of smoothing, with the largest gains for larger smoothing.

**Figure 6:**
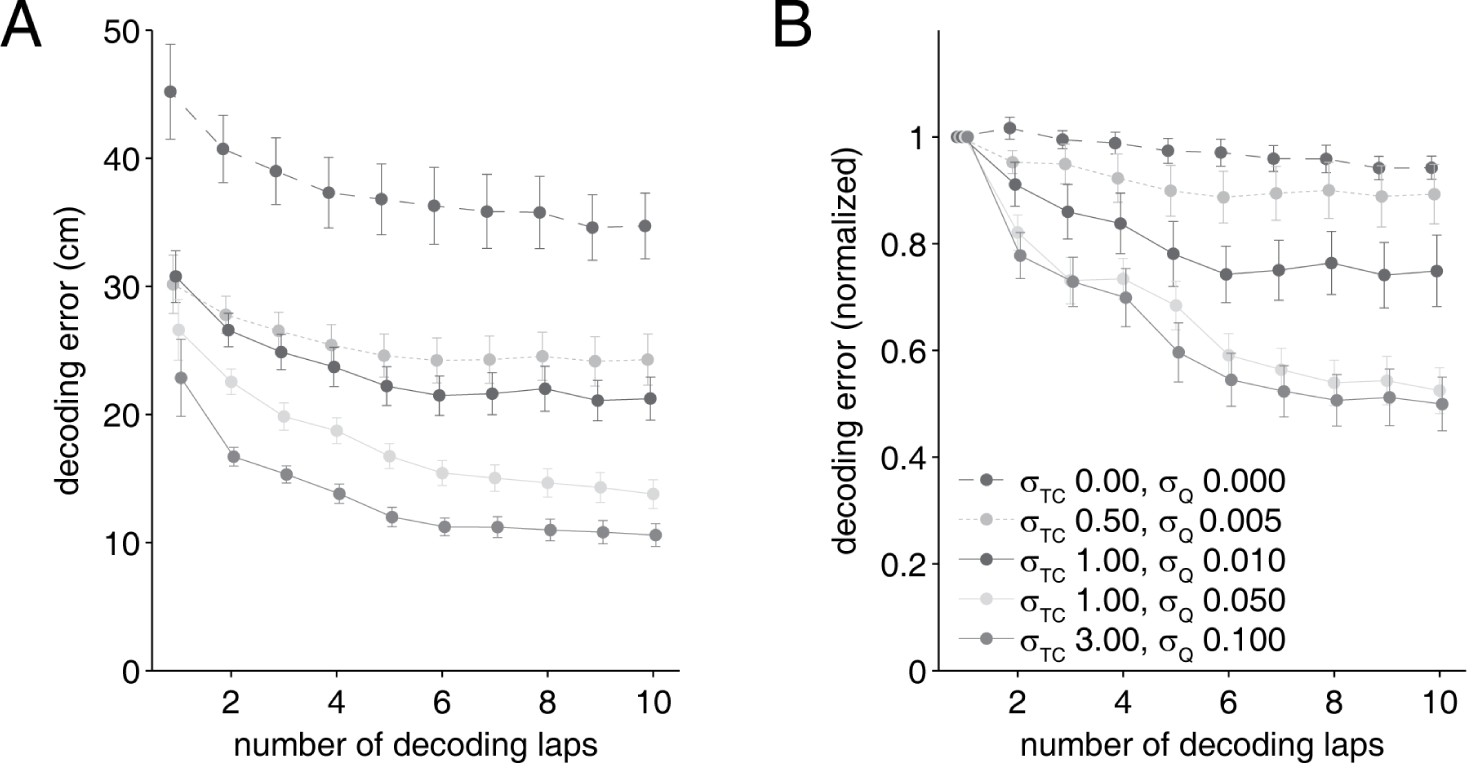
Raw decoding error (**A**) or within-session normalized decoding error (to next-trial decoding error, **B**) as a function of the number of trials included in the cross-validation. Overall, decoding error tended to decrease as more trials were included, but the magnitude of this effect depended on the degree of smoothing used, with stronger smoothing (associated with lower decoding error) benefiting more from including more trials.

The results up to this point raise an obvious question: *why* does decoding performance depend on the way the data is split between encoding and decoding (training and testing) sets? There are two major possibilities.

The first one is overfitting, which assumes that estimating encoding models from a single trial includes fitting a certain amount of noise which generalizes poorly to other trials. In this scenario, including more trials would lead to averaging out of some of this noise, improving performance as shown above (Figure 6). However, a further, non-exclusive possibility is that the encoding model (the mapping between position along the track and neural activity) is not constant across trials. To test this idea, we plotted single-trial decoding performance as a function of the “distance” between the encoding and decoding trials (this can be visualized by shifting the matrix in Figure 3C horizontally, away from its shown configuration with a distance of one trial to distances of multiple trials).

Figure 7 shows that both raw and relative decoding error (normalized within-session to same-trial decoding) tended to increase with larger distance between the encoding and decoding trial (linear mixed model with subject-specific intercepts; effect of trial distance *F* = 10.13, *p* = 0.0017 for parameters with the smallest effect). In other words, decoding was more accurate when using tuning curves estimated from a “near” trial, compared to using tuning curves from a “far” trial. However, it should be noted that pinpointing the source of this effect is challenging, given that aspects of behavior such as average running speed and path stereotypy tend to change over the course of a session, in a manner likely correlated with trial distance (elapsed time) in this experiment. In attempt to control for such changes, we fitted linear mixed models with subject-specific intercepts to the data with decoding error as the dependent variable for each pair of trials (a decoding “target” trial, and an encoding “source” trial). For each such pair we included not only (1) the trial distance (number of trials) and (2) the time difference (between trial start times) as the key regressors of interest, but also the difference in distance run, and (4) the difference in length (in time) between the trials in the pair. Even after the behavioral variables (3) and (4) were included in the model, either trial distance (1) or time difference (2) dramatically improved model fits (nuisance variables only log likelihood -595.2, pseudo-*R*^2^ 0.22; trial distance added -566.39, pseudo-*R*^2^ 0.40, time difference added -563.71, pseudo-*R*^2^ 0.42; all model comparisons *p* < 0.001 for parameters with the smallest effect). This effect suggests that individual trials are associated with distinguishable ensemble firing patterns, potentially reflecting trial-unique aspects of experience (consistent with results from Manns et al. 2007; Mankin et al. 2012; Allen et al. 2012; Ziv et al. 2013). In turn, this observation raises the possibility that a given covert sequence may be best decoded by an encoding model associated with a specific trial.

**Figure 7:**
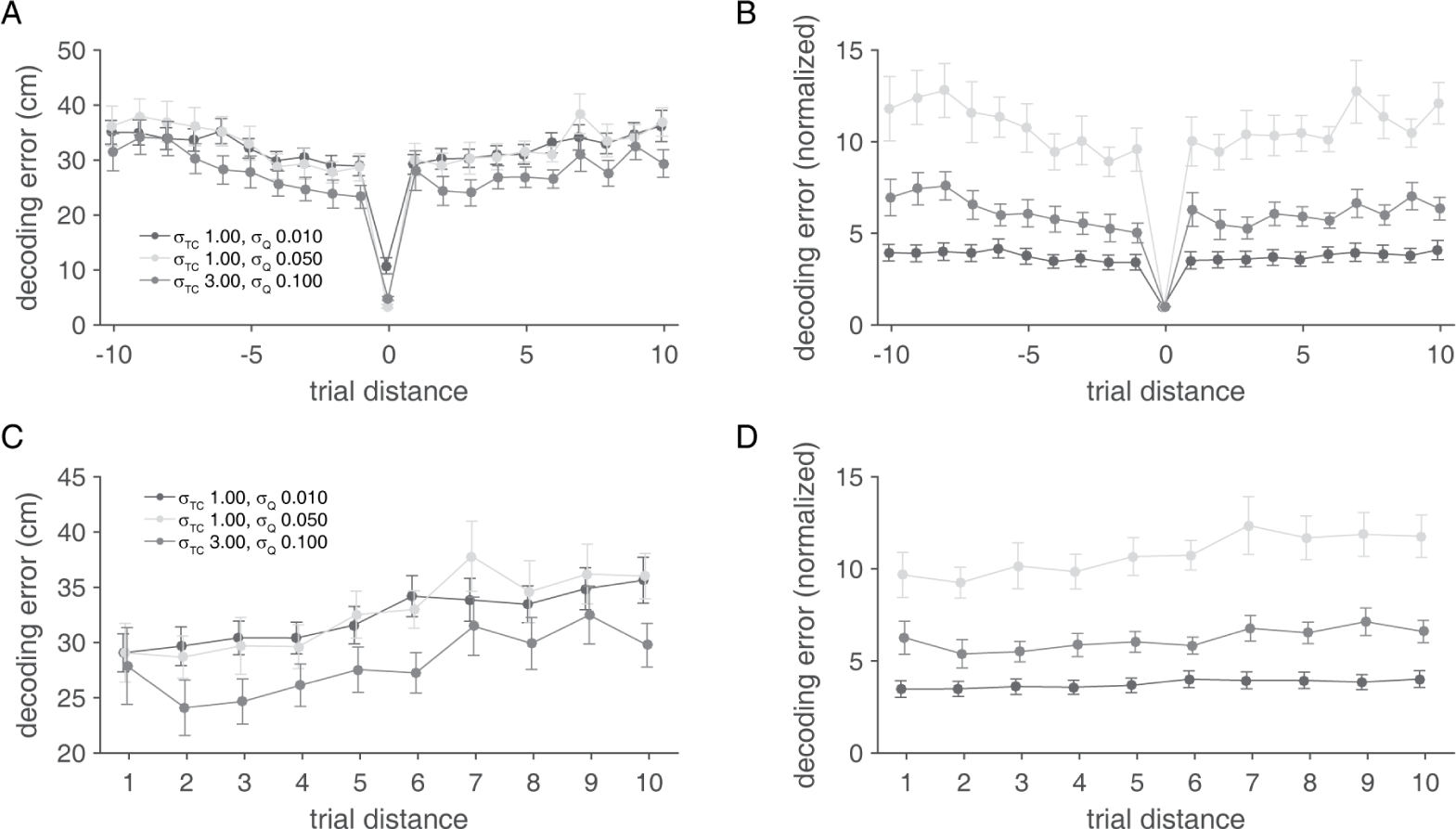
Decoding error as a function of the distance (in number of trials) for single-trials decoding. A trial distance of 0 means that the same trial is used for encoding (estimating tuning curves) and decoding; a trial distance of +1 means that the next trial is used for encoding, and so on. Raw decoding error (**A**) and decoding error normalized within sessions to same-trial decoding error (**B**) tended to increase with larger trial distances. **C** and **D** show the same data but for absolute distance, i.e. previous and next-trial decoding are both distance 1. In order to have sufficient numbers of trial pairs to perform this analysis, trial pairs on which at least 20% of samples could be decoded were included (unlike the 80% threshold used for all other results; see *Methods*).

### Decoding accuracy for different locations

The overall decoding error measure examined so far averages across different stimulus (location) values. However, it is possible that different data splits and decoding parameters differentially affect decoding accuracy for specific locations. Testing whether any such nonuniformity exists in the data is crucial when making comparisons between decoding covert variables (replay) across different stimulus ranges, such as different parts of a track. To test if this occurs, we computed the decoding error as a function of location on the track (Figure 8A-B). Apart from the overall difference in raw decoding error across data splits, there were clear differences in how error was distributed *across* locations: for next-trial and leave-one-out decoding, error tended to increase at the start and end of the track. In contrast, for same-trial decoding, this effect was not apparent at the start of the track. Smaller differences between the same-trial and leave-one-out were also apparent, such as an increase in decoding error around the choice point. Next, we plotted the confusion matrix of actual and decoded locations for the different data splits (Figure 8C). Apart from the overall difference in decoding accuracy, visible as the width of the diagonal, distortions are visible for the leave-one-out case in particular. The point indicated by the white arrow shows relatively poor decoding at the choice point of the T-maze, an effect not apparent for the other data splits.

**Figure 8:**
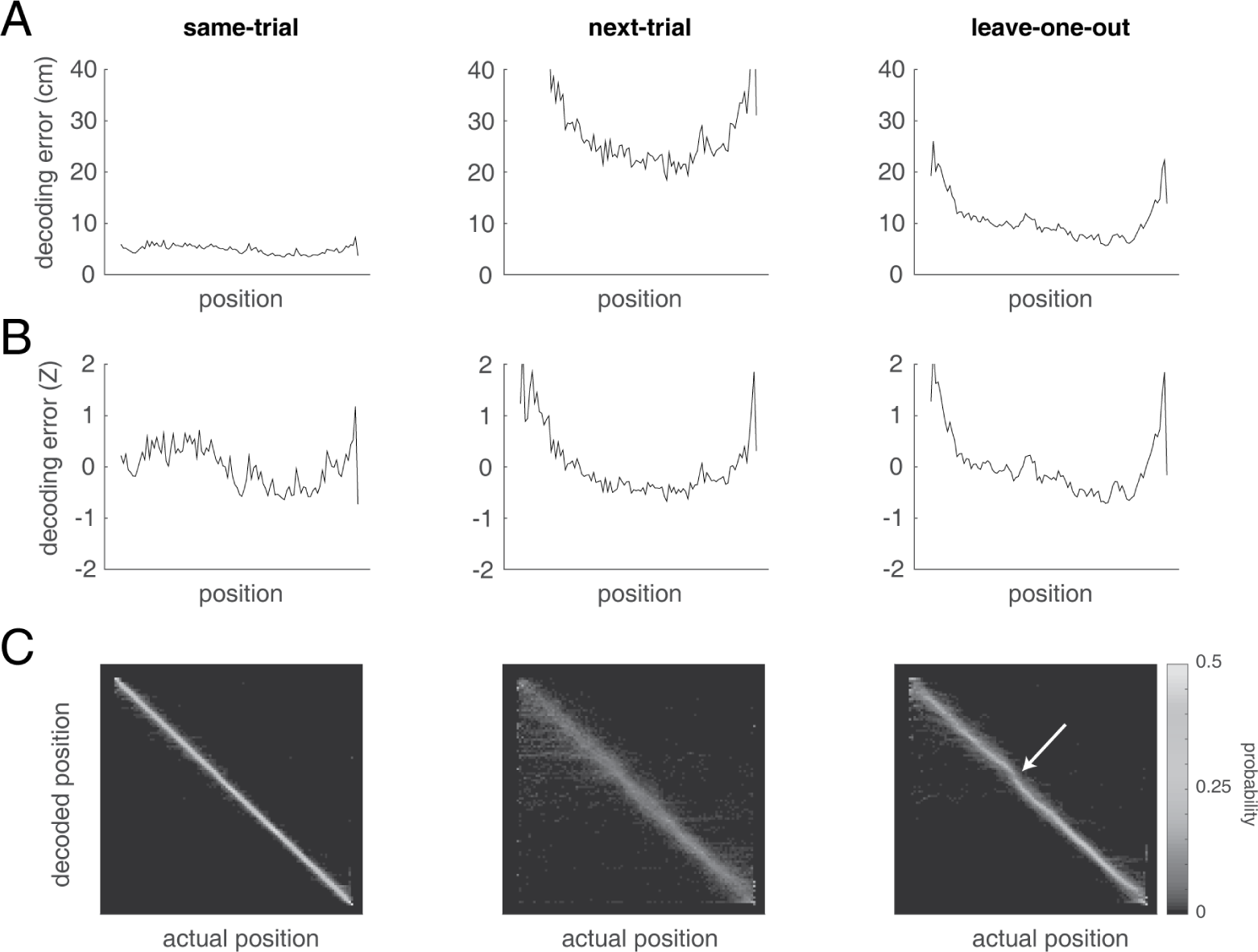
Average decoding error, by location along the track, for the best-performing decoding parameters (starred in Figure 4). Column layout is as in Figure 4), with same-trial decoding on the left, next-trial in the center, and leave-one-out on the right. **A** shows the raw decoding error, **B** shows within-session Z-scored (across space) error. Importantly, these distributions are different; for instance, the next-trial and leave-one-out distributions show increases in error at the start and end of the track not seen in the same-trial distribution. **C**: confusion matrices for actual and decoded position, averaged across sessions. Note the distortion away from the diagonal apparent in the leave-one-out distribution (white arrow) not present in the same-trial case.

In general, there are a number of obvious potential explanations for non-uniform distributions of decoding accuracy, such as differences in the density of place fields and variability in behavior. However, these would be expected to affect both tautological and cross-validated decoding, when the results show strikingly different patterns of decoding accuracy for those cases (Figure 8B). To determine what aspects of the data could help account for the observed nonuniformity in cross-validated decoding, we plotted several quantities related to behavior and neural activity as a function of location (Figure 9). Both average occupancy and its variability across trials were non-uniform (Figure 9A-B) with more sampling around the midpoint of the track compared to the start and end. The average firing rate of neurons with place fields and the number of place fields across the track also showed distributions that did not seem clearly related to decoding accuracy (Figure 9C-D).

**Figure 9:**
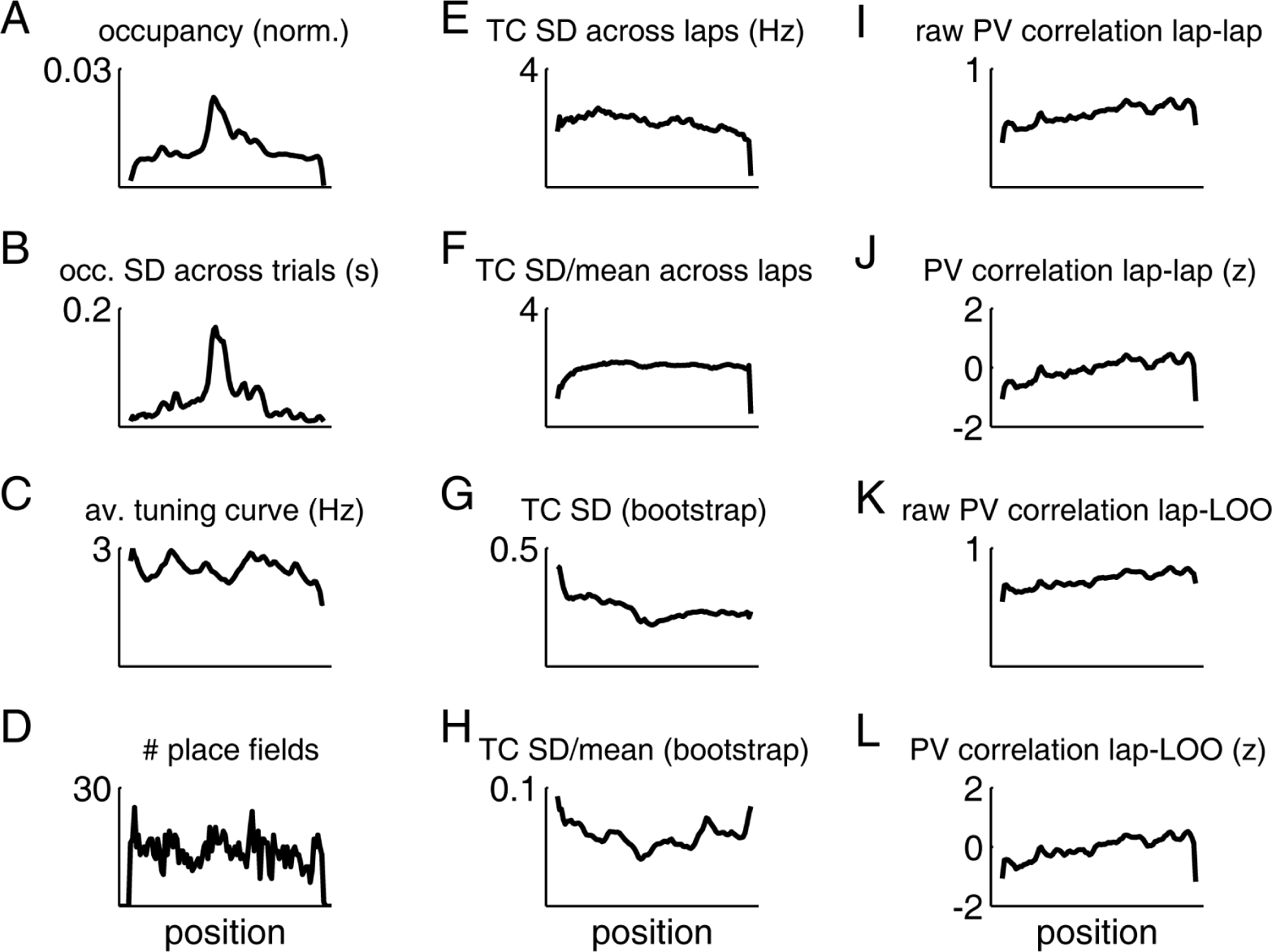
Among several behavioral and neural measures, tuning curve variability as estimated by a bootstrapping procedure is most similar to cross-validated decoding accuracy. **A**: Normalized occupancy (time spent) and its standard deviation across trials within each session (**B**). **C**: Mean firing rate of all cells with at least one place field, and distribution of place field peaks (**D**). **E**: Standard deviation of single-trial tuning curves, computed across trials within a session (raw) or normalized by the mean (**F**). **G**: Entire-session tuning curve variability, estimated by a bootstrapping procedure (raw) or normalized by the mean (**H**). **I**: Raw, and z-scored (**J**) population vector correlation between tuning curves estimated across every pair of trials. **K**: Raw, and z-scored (**L**) population vector correlation between tuning curves estimated from each trial and the complementary leave-one-out tuning curves.

Based on the intuition that cross-validated decoding accuracy depends on the degree of consistency in behavior and neural activity across trials, we computed a number of measures designed to quantify this: (1) the variability in single-trial tuning curves, estimated across trials for each cell individually and then averaged (Figure 9E-F), (2) the variability in entire-session tuning curves, as estimated by a resampling (bootstrap) procedure (Figure 9G-H), and (3) the population vector correlation (i.e. mean firing rates across all cells for a given location), either across single-trial tuning curves or between single-trial tuning curves and the complementary leave-one-out set of tuning curves (Figure 9I-L). These measures showed different distributions across the track, with variability in estimating session tuning curves (Figure 9G-H) showing a distribution most similar to the observed cross-validated decoding accuracy (Figure 8B). Thus, although multiple factors contribute to decoding accuracy, tuning curve variability (as obtained by a bootstrap) may underlie decoding accuracy differences specific to cross-validated decoding.

## Discussion

This study contributes two advances to the methodology for decoding internally generated neural activity. First, we show that using different data splits for the estimation of the encoding model (tuning curves) and the decoding of hippocampal place cell activity affects decoding performance. Specifically, although same-trial decoding was the clear winner in terms of absolute decoding error, single-trial decoding generalizes poorly, leading to large decoding errors when applied to trials other than the one used to obtain tuning curves. Best generalization performance is obtained with leave-one-out cross-validation. These observations are in line with standard practice in the fields of machine learning and statistics (Bishop, 2006); here we explore the implications for the interpretation of covert activity. Specifically, different data splits did not affect decoding performance uniformly across different positions, resulting in biases that need to be taken into account when interpreting decoded “replay” data. The second contribution of this study is that for all data splits, decoding error can be substantially reduced by relatively minimal smoothing, an observation well known in other fields, but not yet systematically applied to hippocampal data.

Both these contributions help address the question of how we should decode and interpret internally generated, covert activity such as occurs in hippocampal “replay” during rest and offline states. The analyses presented here were performed on data from rats running on a T-maze, rather than on covert activity directly. However, the crucial conceptual connection between these two is the following: because the true mapping from neural activity to decoded locations that applies to internally generated activity is typically unknown (see the section below for further discussion), this mapping should be optimized for generalization performance. Operationally, we mimic the decoding of such covert sequences by pretending that we do not know the true encoding model for specific trials on the track – by leaving out these trials in our analysis – essentially treating them as covert sequences, but with the advantage that in this case, we can go back and evaluate decoding performance.

To provide a specific example of how insights obtained from this procedure apply to the interpretation of decoding internally generated activity: suppose we used such decoded locations to detect sequences depicting coherent trajectories along the track. We may find that these “replays” preferentially included the decision point at the middle of the track, rather than the ends of the track. We may be tempted to report this as a finding of interest, perhaps with an interpretation emphasizing prioritized replay as a mechanism useful for reinforcement learning (Schaul et al., 2015; Gershman and Daw, 2017). However, Figure 8 should make it clear that, in the data set used here, such a bias is a straightforward consequence of the increase in cross-validated decoding error at both ends of the track. Crucially, if we had used same-trial decoding error instead, there would not be any indication of a bias favoring the decision point.

Similar to the above example, it is common to use decoding analyses to support a comparison between “replay” of different experimental conditions or spatially distinct areas on the track, such as the left and right arms of a T-maze (Gupta et al., 2010; Bendor and Wilson, 2012; Ólafsdóttir et al., 2015). In such comparisons, it is crucial to ensure that differences of interest in decoded trajectory counts cannot be attributed to intrinsic differences in ability to decode such sequences (e.g. as a result of different distributions of firing fields across locations, firing rates, etc). A common way to control for this is to compare decoding accuracy during behavior across the conditions to be compared; our results show that such measures can differ substantially when based on tautological or cross-validated decoding. Thus, in this setting, as in the previous example, *the cross-validated decoding error provides an important null hypothesis*: the distribution of replay content expected from the decoder’s ability to generalize to neural activity not in the training set.

Note that we are not suggesting any changes to the decoding of “replay” activity itself: this can be done with tuning curves obtained from the full set of behavioral data, because replay activity is not included in the tuning curves. Rather, we point out that the *interpretation* of the replay decoding results should take the cross-validated, not tautological, decoding accuracy during behavior on into account. Whether or not any observed bias in cross-validated decoding error presents a problem depends on the alternative hypothesis to be tested against the potentially non-uniform null hypothesis provided by cross-validated decoding error. Following the example above: the observed bias in cross-validated decoding error to be lower around the choice point of the T-maze casts doubt on the alternative hypothesis that uniform experience is transformed into preferential replay of choice points. However, this same bias may not matter for determining whether there exists a difference between the number of observed “left” and “right” replays.

We found that generalization error depends on the number of trials used to estimate the encoding model, with trial numbers up to the 10 tested generally resulting in lower error. This is intuitive, as a noisy, corrupted tuning curve will lead to a less effective decoder than an accurate one. Note that this implies that when comparing replay content across conditions as in the examples above, the amount of data used to estimate tuning curves should be equalized to eliminate bias due to this effect. As the number of trials used for cross-validation becomes larger, the difference with all-to-all decoding becomes proportionally smaller. Thus, the importance of reporting cross-validated error is especially key when smaller numbers of encoding trials are used, a situation we expect to become more common due to factors such as more complex environments that limit behavioral sampling, and limitations in imaging time due to photobleaching of reporter molecules (Rubin et al., 2015; Malvache et al., 2016).

More trials do not always make for a better encoding model, however; this is illustrated by our observation that decoding error increased when using trials that occurred further apart in time (Figure 7). As we could not explain this effect based on changes in behavioral variables, this suggests a certain amount of trial-unique content, as has been shown previously with different analyses (Manns et al., 2007; Mankin et al., 2012; Ziv et al., 2013). If the contribution of time, or trial-unique features more generally, to internally generated sequences is large (Takahashi, 2015; Schwindel et al., 2016), then averaging across many trials may limit and/or bias the detection of trial-specific replay content. As it is not yet clear to what extent internally generated sequences reflect trial-unique experience, it is difficult to convert this possibility into specific recommendations when interpreting decoded replay data. A conservative approach would be to verify the robustness of a decoding result against variations in the encoding model used (e.g. by using different subsets of trials for decoding; we thank one of the referees for this suggestion).

Finally, beyond the comparison of different data splits discussed above, we show that regardless of split, decoding error can be reduced substantially by decoding spike density functions (SDFs) rather than binned spike counts. Such temporal smoothing has been shown to improve decoding of arm reaching direction from motor cortex activity (Cunningham et al. 2009; see also Kass et al. 2003; Shimazaki and Shinomoto 2010; Prerau and Eden 2011 for a more general treatment of statistical issues in spike rate estimation) but to our knowledge this approach has not been used in studies of hippocampal place cell activity. Although numerous studies have examined the effects of the size of the time window on decoding accuracy (τ e.g. Wilson and McNaughton 1993; Zhang et al. 1998; Resnik et al. 2012; Chen et al. 2016), this is different from our spike density function (SDF) estimation approach: the Gaussian kernel width used in SDF estimation can be manipulated independently from the window size. Thus, for a given window size, such as the 25 ms used here, a variable amount of smoothing can be applied. This modification can be straightforwardly accommodated in commonly used Bayesian decoding procedures (Zhang et al., 1998). Remarkably, decoding performance improves even when estimating SDFs with very narrow kernels (e.g. with a standard deviation of 2 or 5 ms).

Using narrow kernels is particularly important for applications in decoding covert activity, which in the case of hippocampal place cells is temporally compressed relative to behavioral experience (Nadasdy et al., 1999; Lee and Wilson, 2002; Dragoi and Buzsáki, 2006; Buzsáki, 2015).

### Limitations

Our suggestion that decoders intended for covert neural activity should be optimized for cross-validated (generalization) performance is based on the assumption that the “true”, correct decoder for such activity is unknown. Clearly, the approach taken here cannot itself determine the true mapping from covert neural activity to stimulus space. Demonstrating the nature of this mapping is a challenging problem, which may require grounding in experimental observations. Two promising directions may include (1) obtaining access to a brain-internal decoder, such as a downstream projection target, making it possible to determine what aspects of presynaptic activity are distinguished at a next processing stage (Ji and Wilson, 2007; Lansink et al., 2009; Ólafsdóttir et al., 2016; Jadhav et al., 2016); and (2) applying experimental manipulations contingent on decoded content, such that any behavioral effects relative to an appropriate control would constitute evidence that the decoder captures something relevant (clusterless decoding is a promising approach for this; Kloosterman et al. 2014; Deng et al. 2015). A different approach is to construct generative models in an attempt to reproduce experimentally observed activity (Johnson et al., 2008; Pfeiffer and Foster, 2015; Chen et al., 2016). In the limit of a perfect match between the model output and the experimentally observed data, then the optimal decoder can be determined from what is now a known ground truth (the generative model). However, these approaches are not yet mature, and may remain impractical for the purpose of determining the most suitable decoder in any given experiment. Thus, we provide more practical recommendations in the final section.

A different limitation of this study is that although encoding model parameters can be optimized for decoding error when the true location is known, it is unclear how the parameters obtained in this way should be applied to decoding covert activity. Estimates of the temporal compression in internally generated vs. overt activity range from 7-20x (Lee and Wilson, 2002; Davidson et al., 2009; Buzsáki, 2015), thus a practical starting point would be to simply reduce the *σ_Q_* found to be optimal for decoding overt behavior by a factor in that range. For the data set used here this would suggest a value of *σ_Q_* = 5 ms to be a conservative estimate. Future work could provide a more principled estimate of this parameter by, for instance, using generative models as outlined above.

Similarly, our current estimation method for tuning curves uses a relatively *ad hoc* approach of non-parametrically obtaining firing rates from the data, followed by smoothing. Other work has used parametric approaches such as fitting Gaussians or Zernike polynomials (Barbieri et al., 2002); such methods are completely compatible with the approach we take here. Our goal in this study was not to determine which method for tuning curve estimation works best; rather, the main purpose of not using raw, unsmoothed tuning curves here was to prevent inconsistent combinations of spike counts. Looking forward, however, there are clearly opportunities for improving the estimation of tuning curves, such as propagating uncertainty about estimated firing rates throughout the decoding procedure, correcting for the blurring effects of theta phase precession (Lisman and Redish, 2009), and taking the presence of different gamma oscillations into account (Zheng et al., 2016).

Finally, the results provided here are based on one specific data set. However, we emphasize that the specific optimal parameters and decoding error distributions found here are not meant to be imported verbatim to analysis of other data sets for which they may or may not work well; if this was the purpose of the study it would indeed be important to test how consistent the inferred optima are. Rather, these results illustrate the importance of choosing parameters and data splits in a principled manner, and suggests specific steps that can be applied to other data sets to find parameters appropriate for that data.

More generally, although we used hippocampal place cell data from rodents, the ideas developed here can also be applied to other systems in which covert activity can be meaningfully decoded. In rodents, these include the head direction system (Peyrache et al., 2015), areas involved in the processing of decision variables such as orbitofrontal cortex and ventral striatum (Stott and Redish, 2014), and internal representations of time (Pastalkova et al., 2008; MacDonald et al., 2013; Mello et al., 2015). Non-human primate studies prominently explore the generation of motor activity related to upcoming reaching movements (Wu et al., 2006; Yu et al., 2009), and ensemble recording and analysis methods are becoming increasingly common in studies of decision making (Rich and Wallis, 2016). In human subjects, MEG studies have started to explore the fast dynamics of thought (King and Dehaene, 2014; Bellmund et al., 2016; Kurth-Nelson et al., 2016), and MVPA has revealed structure in internally generated activity in a wide range of domains (Reddy et al., 2010; Brown et al., 2016). The present study suggests that the analysis of internally generated sequences of hippocampal activity in rodents can interact productively with statistical approaches developed across domains.

### Summary: three practical guidelines for the decoding and interpretation of internally generated neural activity

The use of cross-validation for decoding is commonplace in human neuroimaging studies (Pereira et al., 2009; Shirer et al., 2012; Varoquaux et al., 2016). Several studies performing position decoding on rodent hippocampus data have used a split between training and testing data (e.g. Zhang et al. 1998; Rutishauser et al. 2006; Davidson et al. 2009; Resnik et al. 2012; Agarwal et al. 2014), but this practice has not been consistently applied in this field. Moreover, the motivation for reporting decoding errors based on cross-validation, and its particular importance for the interpretation of internally generated activity, is typically not made explicit. For the decoding of hippocampal place cell data for this purpose, we suggest the following:

- Report cross-validated, not tautological, decoding error on “running” data. This is good practice in general, but particularly crucial when using decoding accuracy to reveal possible bias in the ability to decode different conditions or trajectories. The cross-validated decoding error distribution should be viewed as a null hypothesis for comparison with alternative hypotheses about the decoded content of internally generated activity, such as replay.
- Because cross-validated decoding error depends (1) on the amount of data (e.g. number of trials) used to estimate tuning curves, and (2) temporal distance between trials used to estimate tuning curves and the time of decoding. Thus, these factors should be equalized, either by design or by subsampling, when comparing replay content across conditions. Even very mild smoothing of the spike trains to be decoded, such as a 5 ms Gaussian kernel for spike density functions, and a 3 cm kernel for tuning curves, can substantially improve decoding performance. However, due to the compression of hippocampal replay relative to behavioral experience, excessive smoothing is discouraged.
- Even very mild smoothing of the spike trains to be decoded, such as a 5 ms Gaussian kernel for spike density functions, and a 3 cm kernel for tuning curves, can substantially improve decoding performance. However, due to the compression of hippocampal replay relative to behavioral experience,excessive smoothing is discouraged.

## Acknowledgments

This work was supported by NWO, HFSP, and the Templeton Foundation (MvdM). We are grateful to Loren Frank, Margaret Carr, and Caleb Kemere for sharing the original design upon which our electrode arrays were based. We thank Nancy Gibson, Jean Flanagan, and Martin Ryan for assistance with animal care, Harmen VanderHeide, Jacek Szubra, Andrew Dubé, and Zhenwhen Zhang for technical assistance, and Min-Ching Kuo for assistance with surgery. We thank Adam Johnson, Elyot Grant and Alireza Soltani for helpful comments on an earlier version of the manuscript, and the referees for insightful suggestions.

## Author contributions

MvdM designed and supervised research. AAC performed experiments, preprocessed and annotated the data. AAC, YT and MvdM wrote analysis code. MvdM performed analysis. MvdM wrote the manuscript with input from AAC and YT.

